# Copy Number Variation-Based Molecular Sexing of *Ixodes scapularis* and *Rhipicephalus microplus* Immature Stages Using qPCR and ddPCR Approaches

**DOI:** 10.64898/2025.12.19.695610

**Authors:** Arvind Sharma, Benjamin Faustino, Isaac Hinne, Jakub Wudarski, Saransh Beniwal, Nurul Hamidi, Kyla Ercit, Sanjay Basu, Malimba Lisulo, Mattia Poletto, Kelly J. Matzen, Monika Gulia-Nuss

**Author notes:** Equal contribution (Arvind Sharma, Benjamin Faustino).

## Abstract

Accurate sex identification of immature ticks is essential for understanding sex-specific ecological dynamics, pathogen transmission, and reproductive biology. However, tick larvae and nymphs lack morphological sexual dimorphisms, limiting the studies. Here, we report the development and validation of copy number variation (CNV)-based molecular sexing approach for two important hard tick species, *Ixodes scapularis,* a major public health vector of human pathogens, and *Rhipicephalus microplus,* a pest responsible for significant economic losses in cattle industry. Using newly published chromosomal-level genome assemblies and whole-genome resequencing data, we identified female-enriched CNVs in two genes, calcium/calmodulin-dependent 3’,5’-cyclic nucleotide phosphodiesterase 1A-like (*NPD*) and rap guanine nucleotide exchange factor 2-like (*RAPGEF2*), and developed SYBR-Green based and probe-based qPCR, and droplet digital PCR (ddPCR) assays using DNA extracted non-destructively from a single leg, preserving ticks for continued feeding, molting, and behavioral analysis. In *I. scapularis*, we were able to assign the sex of 87% of nymphs based on results from at least two molecular tests, and these assignments were confirmed when the nymphs later molted into adults. In *R. microplus*, two qPCR assays revealed clear, sex-linked differences in gene copy number, despite the species having an XX: XO sex system. Together, these findings indicate that CNV-based markers are a reliable and widely applicable method for sex determination in immature ticks.

**Graphical Abstract:** 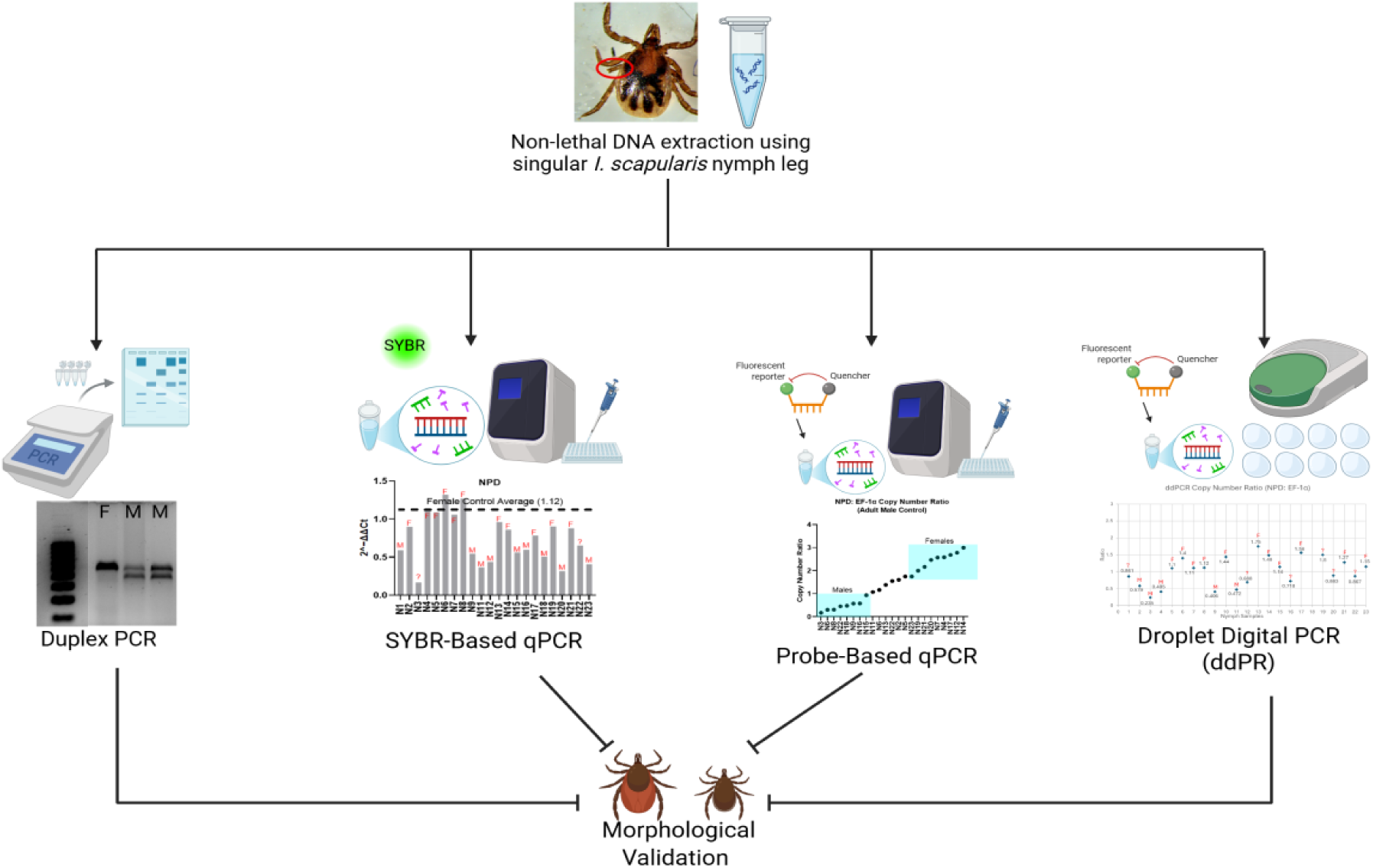

## Background

The increasing prevalence of Lyme disease (LD) and the emergence of other tick-borne human diseases in the United States have become a major public health concern. Ticks are vectors of numerous viruses, bacteria, and protozoa that can impact humans, companion animals, and domestic livestock (Jongejan & Uilenberg, 2004). Blacklegged ticks, *Ixodes scapularis* (Ixodida: Ixodidae), are the primary vector of at least seven pathogens that cause human diseases, including *Borrelia burgdorferi*, the causative agent of LD, *Babesia microti* (human babesiosis), *Anaplasma phagocytophilum* (human granulocytic anaplasmosis), *Borrelia miyamotoi* (hard-tick relapsing fever), *B. mayonii* (new Lyme Borrelia spp.), *Ehrlichia muris*-like agent (ehrlichiosis), and Powassan virus (Powassan virus disease) (Reviewed in Stafford et al., 2017). Since LD was first described in the 1970s, the number of reported human cases in the United States has steadily increased, largely due to the range expansion of *I. scapularis* and spread of *B. burgdorferi* (Diuk-Wasser et al., 2012; Eisen et al., 2016; Kugeler et al., 2015; Pepin et al., 2012). The actual incidence of human disease cases is estimated to be at least 10-fold greater than reported confirmed and suspected cases at ≈450,000 annually (Kugeler et al., 2015).

While *I. scapularis* and LD have been the impetus for recent research on tick ecology and management, the Asian blue tick, *Rhipicephalus microplus*, has a widespread distribution in many tropical and subtropical areas of Asia, South and Central America, and most of sub-Saharan Africa (Angus, 1996; Bermúdez C. et al., 2022; Guglielmone et al., 2006; Kanduma et al., 2020; Tan et al., 2021). It is a one-host tick that primarily infests cattle and is one of the most detrimental vectors of veterinary pathogens, such as *Babesia bovis*, *Babesia bigemina*, and *Anaplasma marginale*, costing the worldwide agriculture industry billions of dollars a year (Bock et al., 2004; Lew-Tabor & Rodriguez Valle, 2016; Todorovic et al., 1981). The spread of *R. microplus* to new areas in the tropics is a significant threat to livestock industries and the livelihoods of rural populations, who often depend on livestock for their survival. The economic importance of *R. microplus* is also compounded by its propensity to displace other native tick species, its higher vectoral capacity, and its ability to develop resistance to acaricides (Abbas et al., 2014; Dzemo et al., 2022; Nyangiwe et al., 2013). Although *R. microplus* has been eradicated from the USA, the threat remains due to cattle exports and the risk of ticks reinfesting from nearby countries through trade (Busch et al., 2014).

The transmission of tick-borne pathogens in natural settings primarily relies on the larval and nymphal stages of ticks, which commonly feed on the same vertebrate reservoir hosts (Piesman & Gern, 2004). Although these early stages are key to pathogen spread, the specific roles of male and female immature ticks remain mostly unknown. One major obstacle in understanding these dynamics is the absence of morphological differences during immature stages. Unlike adults, which can be sexed based on morphological traits, immature ticks are visually indistinguishable, and current sexing methods are often invasive, require specialized equipment, or are only applicable after molting (Dusbábek, 1996; Hu & Rowley, 2000; Keirans et al., 1996).

Molecular sexing methods have been developed for various arthropod species, often utilizing sex-specific DNA sequences to distinguish between males and females. This approach has been successfully applied to organisms such as the silkworm *Bombyx mori* (Abe et al., 1998), the malaria mosquito *Anopheles gambiae* (Krzywinski et al., 2004), and more recently, the black-legged tick *I. scapularis* (Ronai et al., 2025). However, the current tick-specific methods are in the initial stages of development and have not been further adapted into high-sensitivity methods, such as real-time quantitative PCR (qPCR) or droplet digital PCR (ddPCR), which would facilitate broader application in research and colony management. Additionally, the only published method to date is for *I. scapularis* and has not been tested on other tick species.

Ixodidae, the hard ticks, are divided into two subfamilies, Prostriata and Metastriata. The Prostriata contain the genus Ixodes (including *I. scapularis*) and have an XX: XY sex determining system (Gulia-Nuss et al., 2016; Meyer et al., 2010; Nuss et al., 2023). The Metastriata, however, have for the most part lost the Y chromosome and have XX: XO sex-determining system (Chen et al., 1994; Oliver, 1977). *Rhipicephalus microplus* is a metastriate tick with an XX: XO sex-determining system (Newton et al., 1972; Oliver & Bremner, 1968). These differences in sex determination suggest that copy number variation (CNV) in autosomal genes may enable an assay applicable to multiple species. Similar CNV-based approaches have been developed for other arthropods, such as in *Leptinotarsa decemlineata*, the Colorado potato beetle, where sex was successfully identified across different life stages using CNV (Sedláková et al., 2022).

Developing a molecular tool to rapidly and accurately identify male and female ticks at immature life stages is valuable for studying their roles in pathogen transmission and for developing genetic control strategies. Genetic control strategies, including CRISPR-Cas9-based gene drives, control via self-limiting genetic systems (Spinner et al., 2022), and sterile male techniques have been successfully applied in mosquitoes; however, these approaches remain unexplored in ticks (Feng et al., 2021; Kyrou et al., 2018; Wang et al., 2021), primarily due to limited biological and technical knowledge, including the absence of molecular sexing tools. Developing a reliable method to determine tick sex at immature stages will be essential for adapting the genetic control strategies that have proven successful in insect research.

In this study, we developed a molecular sex identification assay using CNVs identified from the recently published chromosomal-level genome assembly of *I. scapularis* and *R. microplus* (Deng et al., 2024; Mou et al., 2025; Nuss et al., 2023; Tidwell et al., 2024). Through bioinformatics approaches, we identified a highly variant gene encoding calcium/calmodulin dependent 3’, 5’ cyclic nucleotide phosphodiesterase 1A like (*NPD*). We compared three approaches for *I. scapularis* sex-determination: SYBR Green-based qPCR, fluorescent probe-based qPCR, and droplet digital PCR (ddPCR) using DNA from a single leg, thus keeping the nymphs intact for feeding. We compared our methods with the published duplex PCR assay for *I. scapularis* (Ronai et al., 2025). After testing with molecular methods, nymphs were blood-fed and allowed to molt, allowing morphological confirmation. We further confirmed *NPD* and an additional CNV gene, rap guanine nucleotide exchange factor 2-like (*RAPGEF2*), in *R. microplus* ticks using a probe-based qPCR assay and whole nymph DNA. This approach offers a flexible and high-confidence method for sexing tick nymphs, a life stage where visual differentiation is not possible, and shows its applicability in more than one species.

## Methods

### Ticks Rearing and Feeding

#### Ixodes scapularis

Pathogen-free nymphs and adults were purchased from the Oklahoma State University (OSU) tick rearing facility and kept in an incubator at 95% relative humidity and 20 °C in our laboratory (Nuss et al., 2017). Nymphs were blood-fed on the dorsal surface of a New Zealand white rabbit (Jackson Laboratories), which was shaved one day before feeding. Based on the putative sex, determined using molecular methods, nymphs were divided into three groups: male, female, and undetermined. Each group was placed in a separate feeding chamber (3-inch plastic capsules) affixed to the shaved skin using Kamar adhesive (Levin et al., 2014; Reyes et al., 2024) and was allowed to feed to repletion. After feeding, individual nymphs were transferred to 1.5 mL tubes and stored at 20 °C and 95% relative humidity for recovery and molting. All procedures were approved by the Institutional Animal Care and Use Committee (IACUC) at the University of Nevada, Reno (IACUC # 21-01-1118).

#### Rhipicephalus microplus

Pathogen-free ticks from the La Minita *R. microplus* strain were sourced from the USDA-ARS (Moscow, ID, USA) and maintained at the Large Animal Research and Imaging Facility, The Roslin Institute (University of Edinburgh). Larvae were blood-fed on female Holstein-Friesian cattle of 3-4 months of age sourced from the University of Edinburgh. Engorged adult female ticks were maintained in 50 mL tubes at 28 °C and 80-90% relative humidity and allowed to lay eggs to recover immature life stages. All procedures were conducted in full compliance with the Animal (Scientific Procedures) Act 1986 (revised 2013). Unfed *R. microplus* adult ticks of known sex, determined by morphological examination, were used to identify variable genes and to develop the sexing assay, and were later included as reference controls to validate the assay. Unfed adults were obtained by removing encased nymphs from the host 12 days post-infestation. Nymphs were allowed to molt at 28 °C and 90% RH for two days. Unfed nymphs were collected five days post-infestation as encased larvae and allowed to molt off host at 28 °C and 90% relative humidity.

### DNA extraction and quantification

#### Ixodes scapularis

The genomic DNA from unfed nymphs and adults was extracted from a single leg by cutting the patella segment of the third leg pair under a dissection microscope. gDNA was extracted using BioSearch QuickExtract™ DNA extraction buffer (Biosearch Technologies #QE09050) following the manufacturer’s protocol. Clipped legs were placed in extraction buffer and vortexed for 15 seconds, followed by incubation at 65 °C for 6 minutes and 98 °C for 2 minutes. The solution was vortexed for 15s between incubations. The concentration of extracted gDNA was estimated by a Nanodrop spectrophotometer and quality of the gDNA was assessed through a PCR amplifying a housekeeping gene (*Actin*). The gDNA was stored at -20 °C.

#### Rhipicephalus microplus

Genomic DNA was extracted using the Qiagen DNeasy Blood and Tissue Kit (Qiagen, Venlo, Netherlands), with the following changes made to the supplementary insect protocol: 1) whole ticks were homogenized with a pestle in 10 μL RL buffer taken from the Norgen Total RNA Purification Kit (Norgen Biotek, Thorold ON, Canada); 2) after the addition of buffer AL and before the addition of ethanol, a second incubation step was added of 10 minutes at 70°C; 3) a second elution step was added where the eluate was pipetted back onto the column and spun again. Eluted DNA was quantified using a NanoPhotometer.

### Gene Selection

#### Ixodes scapularis

Genes with copy number variation (CNV) were identified by comparing normalized read depth from whole-genome resequencing data between male and female *I. scapularis*. To refine this, we calculated coverage across consecutive 2 kb windows and mapped genes to CNV regions using annotations from the GFF (General Feature Format) file.

Eight candidate genes showing sex-specific CNVs were identified from this analysis (Table S1), and all were mapped to the recently published chromosomal-level genome of *I. scapularis* (IscapMGN; GCA_031841145.1).

Primers and fluorescently labeled probes were designed using the *I. scapularis* genome (IscapMGN; GCA_031841145.1) as a reference (Table 1). Candidate gene sequences were selected and assessed for specificity using NCBI Primer-BLAST, and additional off-target screening was performed using NCBI BLAST against the *I. scapularis* genome. Amplicon lengths were kept between 70–150 bp to optimize qPCR performance. Probes were labeled with either 5′-FAM or 5′-HEX fluorophores and a 3′ ZEN Iowa black quencher.

**Table 1:**
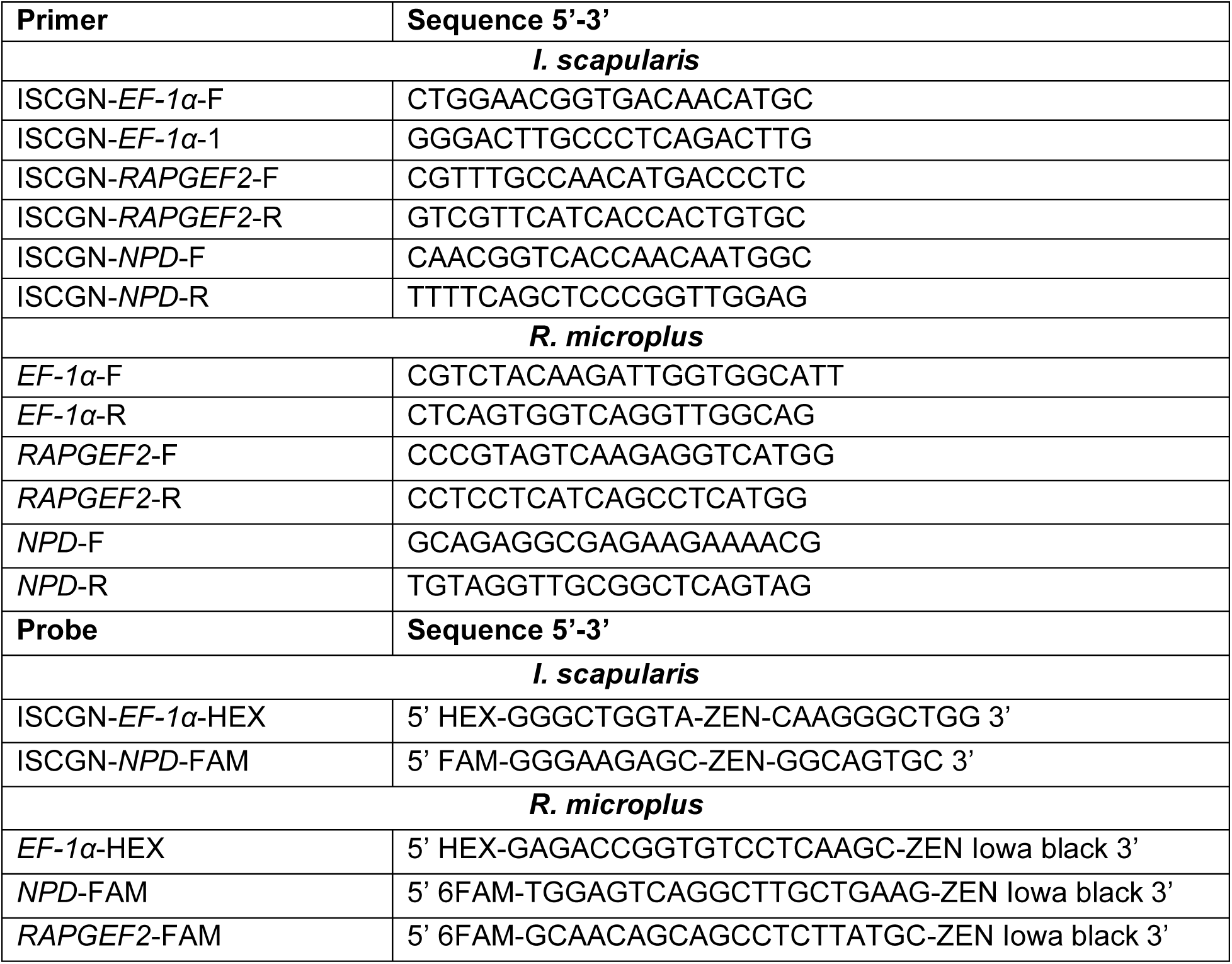
Forward and reverse primer sequences for *I. scapularis* and *R. microplus NPD*, *EF-1α*, and *RAPGEF2*, along with corresponding probe sequences for all three genes in *R. microplus*, used in PCR, qPCR, and ddPCR assays.

Prior to performing any molecular assays, all primer sets were validated using conventional PCR and gel electrophoresis. Each reaction included extracted gDNA from one leg of three separate adult male and female ticks, Taq-Red master mix (Apex #42-137), the respective primer set, and nuclease-free water. PCR cycling conditions are detailed in Table 2, and the PCR products were run on a 1.2% TAE gel.

**Table 2:** PCR, qPCR, and ddPCR cycling conditions, denaturing, annealing, and extension temperatures and time for PCR, qPCR and ddPCR assays.

| **PCR Conditions** |  |  |  |
| --- | --- | --- | --- |
| **Step** | **Temperature** | **Time** | **Cycles** |
| Initial Denaturation | 98°C | 3 minutes | 1 |
| Denaturation | 98°C | 30 seconds | 35 |
| Annealing | 61°C | 50 seconds |  |
| Extension | 72°C | 1 minute |  |
| Final Extension | 72°C | 10 minutes | 1 |
| Hold | 4°C | ∞ |  |
| **qPCR Conditions** |  |  |  |
| **Step** | **Temperature** | **Time** | **Cycles** |
| Initial Denaturation | 95 °C | 10 mins | 1 |
| Denaturation | 95 °C | 11 seconds | 40 |
| Annealing | 60 °C | 30 seconds |  |
| Hold | 12 °C | ∞ |  |
| **ddPCR Conditions** |  |  |  |
| **Step** | **Temperature** | **Time** | **Cycles** |
| Initial Denaturation | 98°C | 3 minutes | 1 |
| Denaturation | 98°C | 30 seconds | 40 |
| Annealing | 60°C | 1 minute |  |
| Extension | 98°C | 10 minutes |  |
| Hold | 4°C | ∞ |  |

#### Rhipicephalus microplus

Two to three candidate genes were selected from each annotated chromosome from the reference genome (BIME_Rmic_1.3: GCF_013339725.1) (Table S2), and primers were designed using the primer design tool in Benchling [Benchling (Biology Software), 2025. Retrieved from https://benchling.com]. Candidate genes were then tested by SYBR Green qPCR to determine their copy number relative to a housekeeping gene known to be present in two copies in each sex, elongation factor 1-alpha 1 (*EF-1α*, LOC119170533). Each 10 μL qPCR reaction contained 5 μL qPCRBIO SYgreen Mix Lo-ROX (PCR Biosystems, London, UK); 0.05 μM of each primer; 5 ng of template gDNA; and molecular biology grade water to make up the rest of the volume. Reactions were run on an AriaMX qPCR system (Agilent, Santa Clara, CA, USA) with the following thermocycling conditions: 3-minute hot start at 95 °C; 40 cycles of (95 °C for 10 seconds, 60 °C for 15 seconds, 72 °C for 15 seconds); melt curve of 95 °C for 3 minutes, 60 °C for 30 seconds, 95 °C for 30 seconds. Candidate genes were evaluated by calculating relative copy number using a -ΔΔCt score (Livak & Schmittgen, 2001) for 19 male and 13 female samples, using the following equation:

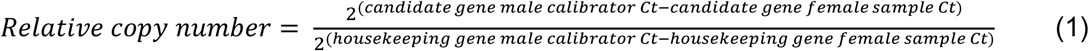

The best candidate genes were identified as those with scores closer to 2 in female samples and closer to 1 in male samples (Table S2). Probe-based qPCR assays were designed using the probe designing tool in Benchling. Candidate gene probes were labelled with 5′-FAM fluorophores, the housekeeping gene *EF-1α* with a 5’HEX fluorophore, and both used 3′ ZEN Iowa black quencher. Final primer and probe sequences for *NPD*, *RAPGEF2*, and *EF-1α* are provided in Table 1.

### Experimental Setup for Molecular Sexing Assays

Using three genes: *NPD*, *RAPGEF2*, and *EF-1α*, identified with CNV in the *I. scapularis* genome, we used three molecular approaches: 1) SYBR Green-based qPCR, 2) fluorescent probe-based qPCR, and 3) ddPCR. Only fluorescent probe-based qPCR was used as a sexing assay in *R. microplus*. We compared these approaches with the published duplex end-point PCR (Ronai et al., 2025).

#### 1) Quantitative PCR (SYBR Green-Based)

DNA from individual *I. scapularis* leg samples was used. Each SYBR-based qPCR reaction was performed in a 10 µL volume containing 0.25 µM of the appropriate primer pair (*NPD/RAPGEF2* and *EF-1α*), 5 µL of SYBR Green Supermix (Bio-Rad; Catalog #1725274), 3.7 µL of nuclease-free water, and 0.5 µL of genomic DNA (∼2.5 ng). All reactions were run in triplicate (CFX Connect™ Real-Time System, BioRad), with each primer set assessed individually. PCR conditions are listed in Table 2. Although both *NPD* and *RAPGEF2* were initially tested, only *NPD* was used in subsequent *I. scapularis* nymph assays due to inconsistent results with *RAPGEF2* primers.

A 2^(-ΔΔCt) method was used to calculate fold change in the target gene expression (Equation 1, Livak & Schmittgen, 2001). Ct values for the target genes were normalized to the housekeeping gene *EF-1α*. Each sample was then compared to a control group, either male or female averages, to determine relative expression. When combining data from multiple plates, an average control fold change value was used to create a consistent reference across all experiments.

#### 2) Probe-based qPCR

For *I. scapularis* probe-based qPCR reactions, the SYBR Green Supermix from the method above was replaced with 5 µL of PCR Biosystems PROBE Master Mix (Batch #240F225K09) and supplemented with 0.25 µM of each gene-specific probe labeled with HEX and FAM fluorophores (Table 1).

For *R. microplus*, two duplex qPCR reactions were performed per sample on an AriaMX qPCR system (Agilent, Santa Clara, CA, USA). Each 10 μL reaction contained 5 μL of pPCR Biosystems Probe Mix Lo-Rox (PCR Biosystems, London, UK); 0.05 μM of each forward and reverse primer (*NPD* or *RAPGEF2* and *EF-1α*); 0.05 μM of each probe; 5 μg of template gDNA; and pure water to make up the rest of the volume. qPCR cycling conditions are outlined in Table 2. For probe-based qPCR assays in both *I. scapularis* and *R. microplus*, copy number was calculated using Equation 1 above.

#### 3) Digital Droplet PCR (ddPCR)

ddPCR was used only for the *NPD* gene, normalized to the housekeeping gene *EF-1α*. Primer and probe sequences were designed based on gene models mapped to the *I. scapularis* Gulia-Nuss genome (Nuss et al., 2023). Adult male and female gDNA samples were included as internal controls in each ddPCR assay to help with the classification of thresholds. ddPCR reactions were performed using the Bio-Rad QX200 Droplet Digital PCR system and ddPCR Supermix for Probes (no dUTP) (Catalog #1863024). Different reaction formulations were assessed:

Each ddPCR reaction was prepared in a total volume of 22 µL, containing 11 µL of 2× ddPCR Supermix, 0.9 µM of each primer, 0.25 µM of each probe, and 2.2–4 µL of genomic DNA (0.5–0.91 ng/µL). The remaining volume was adjusted with nuclease-free water. Reactions were mixed gently before proceeding to droplet generation.

For droplet generation, 20 µL of each prepared reaction mix was loaded into a DG8 cartridge, followed by the addition of 70 µL of droplet generation oil. After droplet formation using the Bio-Rad AutoDG Droplet Generator, 40 µL of the generated droplet (sample) was transferred to a 96-well PCR plate for amplification. PCR cycling conditions are outlined in Table 2.

Following amplification, droplets were read using the QX200 Droplet Reader. Data were analyzed with Bio-Rad QX Manager™ software, and target-to-reference copy number ratios were calculated from the absolute quantification output. These ratios, along with the data from the other molecular methods, were used to classify the 23 nymphal samples as male, female, or unknown based on relative enrichment of the *NPD* target.

#### 4) Duplex PCR

This method uses two primer pairs in a single reaction: one amplifies a 326-bp male-specific region, and the other amplifies a 406-bp region of the autosomal 40S ribosomal protein S4-like gene. *I. scapularis* nymph samples showing both the 326-bp and 406-bp bands were identified as male, while those with only the 406-bp band were identified as female, as described by Ronai et al., (2025). The test samples consisted of nymphal leg DNA, and two adult males and one adult female were included as positive controls. A no-template reaction was included as a negative control. Details on cycling conditions and additional methods are provided in Ronai et al., (2025).

## RESULTS

### Sex determination assays for *Ixodes scapularis* nymphs

#### Gene selection

In *I. scapularis*, eight candidate CNV genes were identified (Table S1), using the published chromosomal-level genome (IscapMGN; GCA_031841145.1) (Nuss et al., 2023). Out of the top eight, calcium/calmodulin-dependent 3’,5’-cyclic nucleotide phosphodiesterase 1A-like (*NPD*, ISCGN302940) and rap guanine nucleotide exchange factor 2-like (*RAPGEF2*, ISCGN121770) were selected as target genes and for primer design as they displayed elevated copy numbers in females across two HiFi datasets: Female-1 (∼2,000 kb) and Female-2 (∼1,800 kb). These two candidate genes are highlighted in Table S1, where ’Start’ and ’End’ indicate coordinates of variation, ’Window’ reflects the number of 1 kb segments, and ’Fold Change’ shows copy number differences between sexes, summarized as male- or female-enriched. Elongation factor 1-alpha (*EF-1α*) was identified as a reference (housekeeping) gene due to its consistent expression across sexes in other tick species, serving as a normalization control (Kim et al., 2023; Salata et al., 2020).

In *R. microplus*, CNV genes were identified on chromosomes 2 and 8. Similar to *I. scapularis*, the most consistently variable CNV genes were identified as *NPD* (LOC119176070) and *RAPGEF2* (LOC119161094) (Table S2).

#### Validation of Sex-Specific Primers and DNA Extraction Efficiency Using Conventional PCR

For *I. scapularis, RAPGEF2*, *NPD*, and *EF-1α* genes were used for conventional PCR to test the amplification from gDNA extracted from the legs of male and female adults. All amplicons were detected at ∼100-125 bp, as expected from the sequence, with minimal to no non-specific bands (Fig S1). While some reactions showed lower expression, the overall results confirmed that the DNA extraction protocol using a single tick leg was effective and that the primers were gene specific (Fig S1).

##### Nymphal DNA

For all the following assays, DNA extracted from a single leg of an individual nymph (N=23) was used. Out of 23 nymphs, three DNA samples failed to amplify with all four assays (Table 3; however, the DNA quality appeared good when a PCR was run with β-actin and run on the gel (Fig S2). Therefore, although these samples did not produce results in our assays, we have included them in the analysis.

**Table 3:** Comparison of sex classification results across molecular assays. This table summarizes the concordance of sex classifications obtained from four molecular methods: Duplex PCR, SYBR Green qPCR, probe-based qPCR, and droplet digital PCR (ddPCR). Nymphal samples (N1–23) are grouped by putative sex based on molecular data, with “M” indicating male and “F” indicating female. Only two nymphs showed full agreement across all four methods. However, another eight nymphs had matching results from three molecular methods, and another ten had matching results from at least two assays. Samples with conflicting or incomplete data are noted as unclassified (-). Samples were sexed if results from at least two molecular methods agreed. Three nymphs, marked with an asterisk (*), died before initiating feeding. The gray rows represent samples that could not be amplified despite having good DNA quality (Fig S2).

| Nymph Sample | Duplex PCR | Probe-based qPCR | SYBR-based qPCR | ddPCR | Assigned Sex | # of Matching Molecular Methods |
| --- | --- | --- | --- | --- | --- | --- |
| N1 | M | - | M | - | M | 2 |
| N2 | - | - | F | M | ? | 0 |
| N3 | M | M | - | M | M | 3 |
| N4 | - | F | F | M | F | 2 |
| N5 | - | F | F | F | F | 3 |
| N6 | - | M | F | F | F | 2 |
| N7 | - | F | F | F | F | 3 |
| N8 | - | M | F | F | F | 2 |
| N9* | M | M | M | M | M | 4 |
| N10 | - | - | - | F | ? | 0 |
| N11 | M | M | M | M | M | 4 |
| N12 | M | F | M | - | M | 2 |
| N13 | - | - | F | F | F | 2 |
| N14 | - | F | F | F | F | 3 |
| N15 | - | M | M | F | M | 2 |
| N16 | M | M | M | - | M | 3 |
| N17* | M | F | F | F | F | 3 |
| N18 | F | M | M | - | M | 2 |
| N19 | - | F | F | F | F | 3 |
| N20 | M | F | M | - | M | 2 |
| N21* | - | F | F | F | F | 3 |
| N22 | - | M | - | - | ? | 0 |
| N23 | - | F | M | F | F | 2 |

#### Duplex PCR

The duplex PCR assay generated correct bands for nine of the twenty-three *I. scapularis* nymphal samples. Of these, eight displayed two bands corresponding to the male-specific 326-bp fragment and the 406-bp autosomal fragment, indicating male sex, while only one sample showed only the 406-bp autosomal band, indicating female sex (Fig S3, Table 3). While some of the samples produced amplification, many bands were faint and difficult to interpret confidently, likely due to the limited gDNA available from a single nymph’s leg. Additionally, using higher gDNA amounts in the duplex PCR led to non-specific bands, which made determining the sex even more challenging, an observation similar to Koloski et al., (2025). This limitation prompted us to explore more sensitive and quantitative molecular approaches to improve accuracy and reliability in sex determination.

#### SYBR-based qPCR

To evaluate the SYBR Green qPCR approach, we initially tested nine each of adult males and females to establish a baseline amplification and confirm the reliability of the method before applying it to immature life stages. This step was essential to ensure the accuracy of our results when testing nymph samples. This method was then applied to 23 nymphal samples for further validation.

The assay for adult *I. scapularis* samples yielded a fold change (2^-ΔΔCt) for males ranging from 0.39 to 0.79 when using *RAPGEF2* as the gene of interest and 0.13 to 0.48 when using *NPD relative to EFα.* For both target genes, all male samples showed lower fold change values compared to the average of the female controls (Fig 1 A, B). This matched the CNV identified in the *I. scapularis* genome, which showed both genes of interest being enriched in the females (Table S2). Following validation with adult *I. scapularis* samples, both SYBR and probe-based qPCR methods were applied to nymphs. Adult male and female gDNA were included in both methods as positive controls to determine thresholds and to establish baseline amplification patterns for probe-based qPCR.

**Figure 1:**
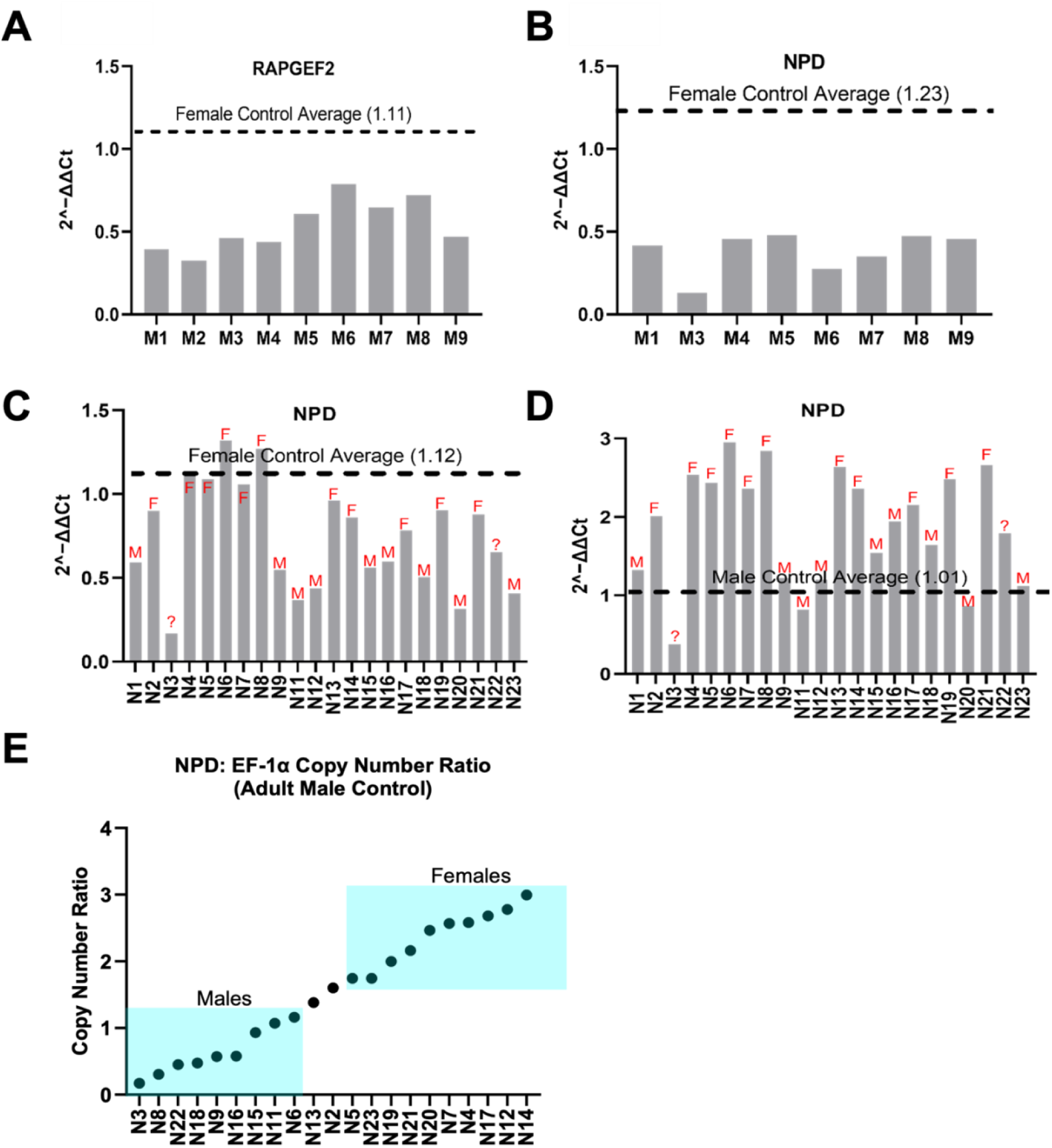
Comparison of fold change variation between adult male and female genomic DNA samples. Relative fold change levels were calculated using the 2^−ΔΔCt method, comparing male and female samples. **(A)** *RAPGEF2* was used as the target gene, with *EF-1α* as the housekeeping gene. The female sample serves as the control reference, normalized to a fold change of 1.11. All nine male samples exhibit a decreased fold change relative to the female average. **(B)** The same analysis was conducted using *NPD* as the target gene, with *EF-1α* again serving as the housekeeping gene. The female control is normalized to 1.23. All 9 male samples displayed a fold change below the female average. However, male 2 from this data set was excluded due to outlier fold change values with *NPD* as the target gene (0.914). **(C) SYBR-based qPCR assay to determine sex of *Ixodes scapularis* nymphs**. DNA from a single leg collected individually from 23 *I. scapularis* nymphs was used for qPCR Relative fold change levels were calculated using the 2^−ΔΔCt method. *NPD* was used as the target gene and *EF-1α* as the housekeeping gene. Adult female samples serve as the control reference, normalized to a fold change of 1.12. **(D)** *NPD* was again used as the target gene, with *EF-1α* serving as the housekeeping gene. Adult male samples serve as the control reference, normalized to a fold change of 1.01. Nymph sample three could be classified as male when using adult female samples as the control; however, it did not align with expected patterns when adult males were used as the control, resulting in inconclusive data. Based on the overall results, 9 samples were classified as male, 11 as female, and the remaining samples were considered undetermined. **(E)** Probe-based qPCR assay to determine sex of *Ixodes scapularis* nymphs. DNA from a single leg collected individually from 23 *I. scapularis* nymphs were used for qPCR. Each point represents an individual nymph sample, with copy number values calculated relative to the reference gene *EF-1α*. Thresholds were established using adult male and female samples to interpret copy number results: samples with an *NPD*: *EF-1α* ratio ≤1.16 was classified as male, those ≥1.71 as female, and values between these thresholds were considered undetermined. Using this approach, 19 samples were confidently classified, 9 as male and 10 as female. Two samples (N1 and N10) produced unusually high copy number values (above 5) and were excluded from the analysis as outliers.

Of the two target genes initially tested on *I. scapularis* nymphs, *RAPGEF2* yielded contrasting results and was therefore excluded from further analysis, and only *NPD* was used (Fig S4). The qPCR results of nymphs showed fold change patterns similar to those previously observed in adults. When adult female samples were used as the control, nymphs predicted to be males showed a low copy number in *NPD* (fold change <1), while nymphs predicted to be females showed higher copy numbers ∼1 (Fig 1C). This pattern was reversed when adult males were used as the control, with female nymphs exhibiting a clear increase in *NPD* fold change (Fig 1D). Threshold values were established using adult male and female controls: 1.12 when females were used as the reference and 1.01 when males were used. These thresholds provided a benchmark for classifying unknown samples. Using this approach alone, we could confidently predict the sex of 20 out of 23 nymphs (87%) (Table 3). Three samples failed to amplify; therefore, this could be considered 100% if the failed samples are removed.

#### Probe-Based qPCR

For probe-based qPCR in *I. scapularis*, copy number was assessed using the *NPD*: *EF1α* copy number ratio. Samples with ratios ≤ 1.16 were interpreted as male, while ratios ≥ 1.71 were considered female. These threshold values were based on average ratios obtained from male and female positive control samples. This method allowed us to classify 19 out of 23 samples (82.6%), and all but 5 matched the classifications (from those available) obtained from SYBR Green-based qPCR data (Fig 1E, Table 3). These results further support *NPD* as a reliable target for molecular sexing of *I. scapularis* nymphs.

#### ddPCR

To further validate the sex of the *I. scapularis* nymphal samples, ddPCR was used to measure *NPD* ratio levels, normalized to the *EF-1α* reference gene. Adult male and female gDNA were included in each run as internal controls to help set classification thresholds. Different reaction conditions were evaluated, and variability between replicates was reduced by increasing the gDNA input. Initial reactions (22 µL) used 1 µL genomic DNA (0.23 ng/µL) with standard primer (0.9 μM) and probe (0.25 μM) concentrations but showed high variability and large error bars (Fig S5). To improve precision, we optimized the assay by increasing genomic DNA input to 2.2 µL and 4 µL (0.51–0.92 ng/µL) and adjusted primer and probe concentrations accordingly. The 2.2 µL input followed Bio-Rad’s recommended protocol. All results were generated using these optimized conditions.

A copy number ratio generated from the Bio-Rad QX Manager™ software of ∼0.6 or lower was classified as male, while ratios ≥1.0 were considered female, based on consistent patterns observed in adult controls (Fig 2A). Using this method, we were able to confidently classify 17 of the 23 nymph (74%) samples as male or female (Fig 2B). Of these, 13 samples matched classifications from at least one of the SYBR or probe-based qPCR methods, supporting the reliability molecular tools for sex determination in *I. scapularis* nymphs (Table 3).

**Figure 2:**
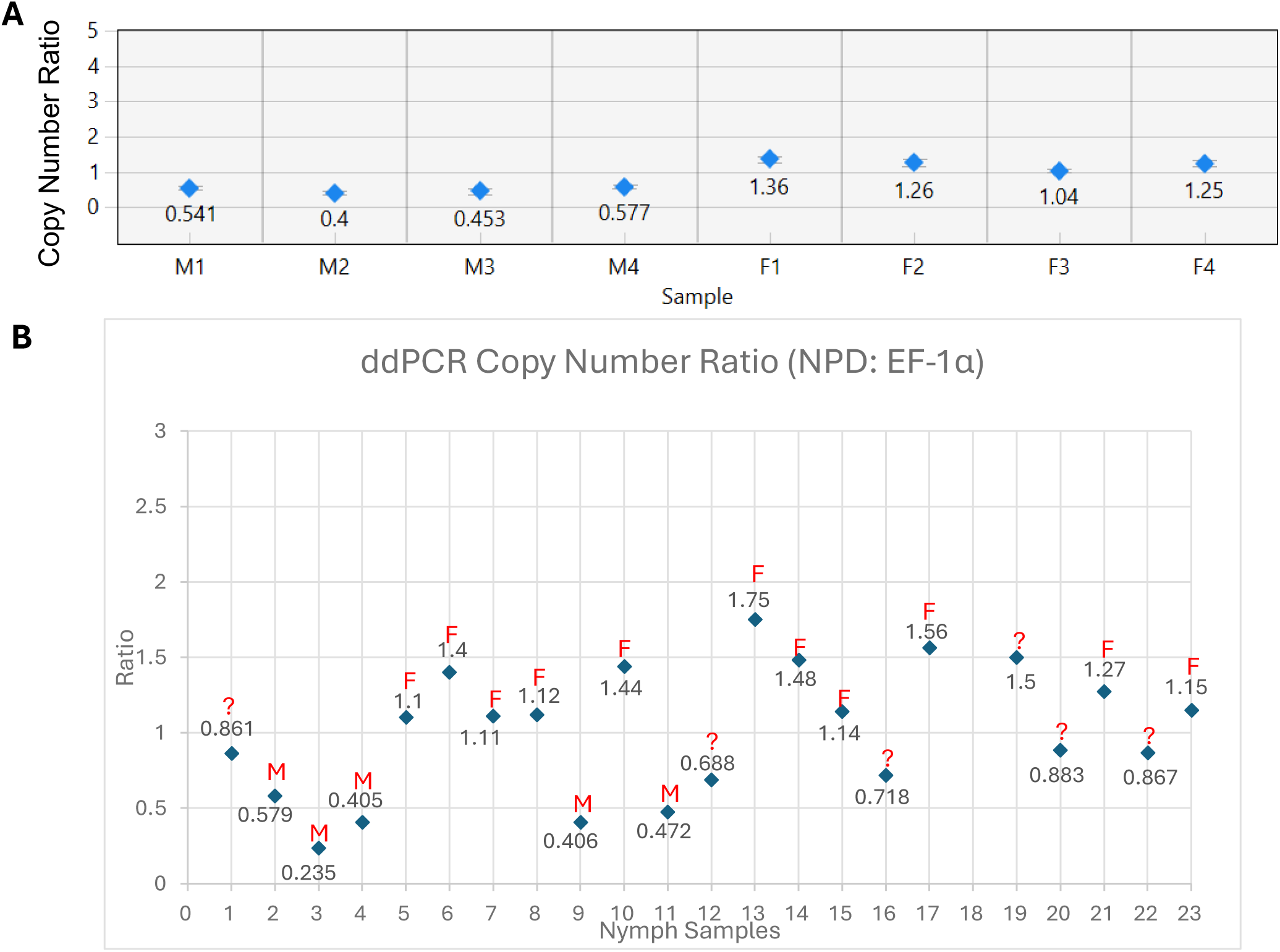
ddPCR assay to determine sex *of Ixodes scapularis.* **(A)** Adult control samples (4 males and 4 females) were used to establish thresholds. Males consistently showed *NPD*: *EF-1α* ratios ≤0.6, while females showed ratios ≥1.0. These clear and reproducible results provided the basis for classifying nymph samples. **(B)** Nymphal samples were analyzed using 2.2-4 µL (0.5ng/uL-0.91ng/uL) of gDNA per reaction. Based on the thresholds established from adult controls (males; ≤0.6 and females: ≥1.0), 17 of the 23 samples were confidently classified as male or female, with 5 being classified as male, and 12 being classified as female.

#### Comparison of Molecular Sexing Methods and Morphological Validation of I. scapularis Nymphs

To assess the accuracy and consistency of our molecular assays, we compared results from all four molecular methods (Duplex PCR, SYBR Green qPCR, probe-based qPCR, and ddPCR. Samples were grouped by putative sex: male (M), female (F), or undetermined (?), based on the molecular data (Table 3). Sex was assigned when at least two molecular methods produced consistent results. For example, if both SYBR and ddPCR identified a sample as female while the probe-based method failed or gave conflicting results, the sample was classified as female. This consensus-based approach helped reduce the impact of technical variability from any single method.

Of the 23 nymph samples tested, we were able to assign sex to 20 individuals (87%), nine males and 11 females, based on consistent results from at least two molecular methods. Two nymphs showed complete agreement across all four assays. An additional eight nymphs had matching results from three methods, and 10 more showed agreement between two. The remaining three samples could not be confidently classified due to conflicting results or failed amplification (Table 3). To rule out DNA degradation as a cause of failed amplification, all nymphs were tested for the actin housekeeping gene, which consistently amplified in the three unclassified samples, indicating good DNA quality (Fig S2). Therefore, the lack of sex assignment for these samples remains unexplained and may be due to other technical or biological factors such as the presence of PCR inhibitors.

#### Morphological validation of Sex

Of the initial 23 nymphs, 20 survived until ready to feed (∼6 weeks), 3 died prior to feeding. Of these three, 1 was classified as female and 2 as males, according to results from 4 assays (Table 3). Of the 20 surviving nymphs, 10 fed to repletion and 7 molted successfully, allowing for morphological verification of sex and evaluation of assay accuracy (Fig 3 A, B).

**Figure 3:**
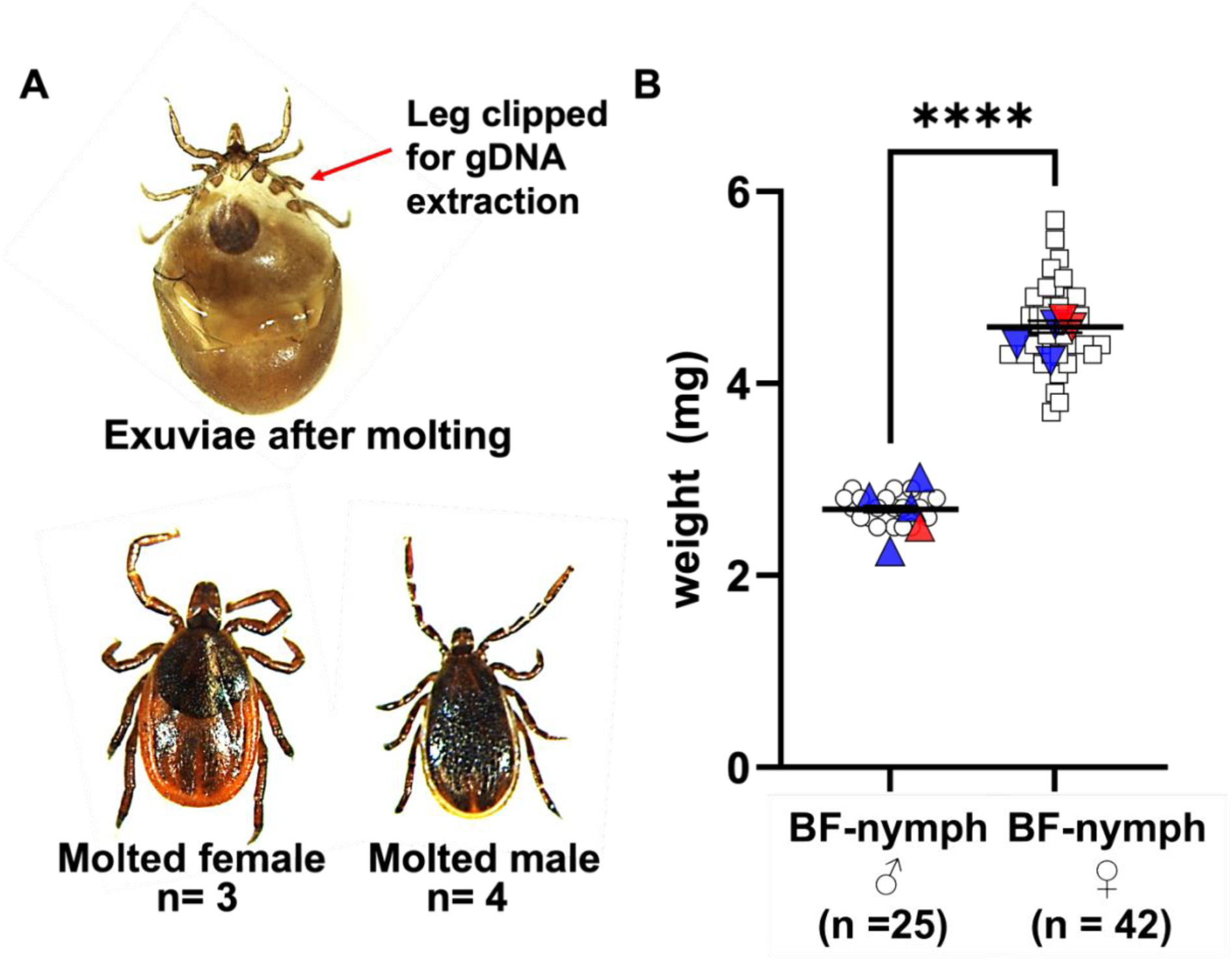
Adult *Ixodes scapularis* molted from the molecularly sexed nymphs. **(A)** Exuviae (molted casing) left behind after an *I. scapularis* nymph molted to the adult stage. The arrow indicates the location where a single leg was clipped for non-lethal DNA extraction; the missing leg regenerated in the resulting adult. Representative image of adult male (right) and adult female (left) that successfully molted from nymphs whose sex was determined using the CNV-based molecular assays. **(B) Weight of blood-fed *I. scapularis* nymphs.** Fifty-seven blood-fed nymphs were weighed to evaluate body mass as a predictor of sex, resulting in provisional classification of 20 males and 37 females. The body weights of 10 nymphs previously identified as male (n = 4), female (n = 3), or undetermined (n = 3) by molecular sexing fell within the same corresponding weight categories. Molecularly confirmed males (blue triangles) and females (blue inverted triangles) are shown in their respective groups. Nymphs that could not be sexed molecularly were assigned to male or female categories based solely on body weight (red triangles and red inverted triangles). Statistical significance was calculated by unpaired t-test with Welch’s correction was used for comparing control using GraphPad Prism v10. **** p < 0.0001.

For nymphs predicted to be male by molecular methods, 3 of 4 (75%) that survived to molting emerged as adult males. One assigned male nymph died after 2 months post blood meal, before molting. For those predicted to be female, 2 of 3 (66.6%) individuals molted into an adult female, 1 of them died after blood meal. Despite the small number of individuals that molted, these results provide preliminary morphological support for the molecular sex classification.

#### Nymphal Body Weight as a Predictor of sex

We also tested whether body weight could be used to predict sex. Fifty-seven blood-fed nymphs were weighed to evaluate body mass as a predictor of sex. resulting in provisional classification 20 nymphs had an average weight of 2.7 mg and 37 nymphs weighed on an average 4.4 mg, and were classified as males and females, respectively (Fig 3C). The body weights of seven nymphs previously identified as male (n = 4) and female (n = 3) by molecular sexing fell within the same corresponding weight categories. Nymphs that could not be sexed molecularly (undetermined, n=3) were assigned to male or female categories based on body weight. These 3 unclassified nymphs fell within these ranges, allowing us to designate 2 as likely female and 1 as likely male. Of these, 1 weight-assigned female died after feeding (∼2 months), but the other individuals survived to molt and confirmed the predicted sex, supporting the utility of weight as an additional indicator.

### Sex determination assay for *Rhipicephalus microplus* nymphs

#### Probe-Based qPCR

In *R. microplus*, two dual probe-based assays were similarly designed to target *RAPGEF2* and *NPD* respectively, with *EF-1α* as the housekeeping gene. These assays were validated and optimized using sexed adult male and females. The mean relative copy number of *NPD* was 0.99 in males and 2.02 in females, and of *RAPGEF2* was 0.81 in males and 1.41 in females (Fig 4A). While both the *NPD* and *RAPGEF2* assays individually produced reliable results, using them together helped boost confidence in the findings.

**Figure 4:**
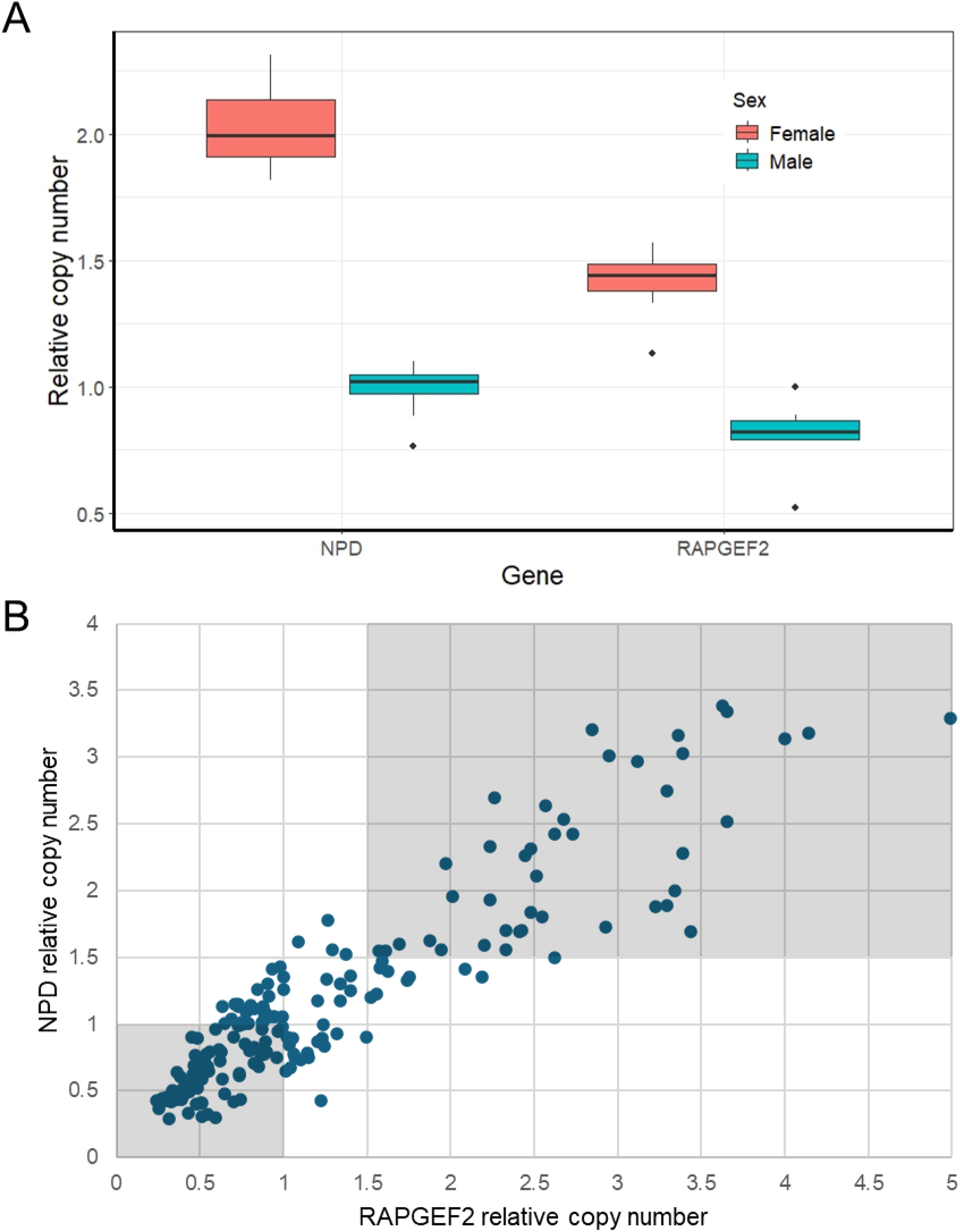
Probe based qPCR assay to determine sex *of R. microplus*. **(A)** Boxplot of relative copy number of genes *NPD* and *RAPGEF2* in male and female adult *Rhipicephalus microplus*. Values were obtained from a dual probe qPCR assay using *EF-1α* as a housekeeping gene. **(B)** Results of dual probe-based qPCR assay on unsexed *Rhipicephalus microplus* nymphs. Both *NPD* and *RAPGEF2* gene assays were used to sex 201 nymphs. Each point represents an individual nymph sample, with copy number values of *NPD* or *RAPGEF2* calculated relative to *EF-1α*. Thresholds were established using adult male and female samples to interpret copy number results. Samples with relative *NPD* and *RAPGEF2* copy number under 1 were classified as male and above 1.5 were female. Values between these thresholds were considered undetermined.

When using this assay to sex nymphs, we used a more conservative window of acceptable values, as samples returned a wider range of copy number values than adult controls. Any sample which produced a Ct value of 32 or above was omitted from the assay as was assumed to have low template quality. Samples with a relative copy number of above 0.5 and below 0.2 were also omitted from sexing. Samples with both relative *NPD* or *RGNEF* copy number below 1 were considered to be male and above 1.5 were considered to be female. Determination of the sex of individuals displaying intermediate copy number values was determined unreliable (Fig 4B). Out of 201 nymphs tested with these assays, six samples were discounted due to high Ct values. Nine samples were discounted due to too high or too low relative copy numbers. Of the remaining samples, 78 nymphs were assessed to be male, and 42 were female. The remaining 66 nymphs had intermediate copy numbers which could not be unequivocally assigned to either sex. Copy number results from the *NPD* assay fell into a more bimodal distribution than *RAPGEF2* (Fig S6), and there was a larger gap in mean copy number among adult samples in the *NPD* assay. Therefore, the *NPD* single assay may be reliable enough to be used on its own.

## Discussion

This study highlights the development of molecular sex identification methods for *I. scapularis* and *R. microplus* nymphs using a female-enriched gene, calcium/calmodulin-dependent 3’,5’-cyclic nucleotide phosphodiesterase 1A-like (*NPD*), detected via SYBR Green qPCR, probe-based qPCR, and droplet digital PCR (ddPCR).As shown for *I. scapularis*, these approaches require only a small amount of tissue, such as a single leg or a portion of a leg, making it possible to determine sex without sacrificing the tick, allowing for continued monitoring or downstream validation. To our knowledge, this is the first adaptation of high-sensitivity molecular sexing assays such as qPCR and ddPCR for immature stages in any tick species, addressing a critical gap since sex can typically not be distinguished morphologically until after molting to adults.

Accurate and early sex identification of ticks is important for understanding population sex ratios. Early sex identification also enables researchers to study how male and female ticks differ in traits like host-seeking, blood-feeding, and pathogen acquisition factors that directly influence their role in disease ecology (Kiszewski et al., 2001; Lefcort & Durden, 1996; Troughton & Levin, 2007; Varone et al., 2025). Additionally, early sexing is essential for applying genetic control strategies involving self-sexing strains, which have been successful in mosquitoes but remain underdeveloped in ticks (Feng et al., 2021; Kyrou et al., 2018; Spinner et al., 2022; Wang et al., 2021).

Previous efforts to sex immature ticks often rely on destructive dissection, high-end microscopy, or genetic markers not yet validated in qPCR-based platforms (Hu & Rowley, 2000; Keirans et al., 1996). Recently, Ronai et al. (2025) developed a duplex PCR-based method for molecular sexing in *I. scapularis*, which targets both a male-specific and an autosomal fragment, enabling sex determination in immature stages. However, when we applied this method using non-lethal DNA extraction from a single leg, the resulting bands were faint and inconclusive (Fig S3). Attempts to optimize the protocol by increasing either the amount of gDNA or the number of PCR cycles led to the appearance of spurious bands, ultimately making the method unreliable and inconsistent for our non-lethal sampling approach (our data and Koloski et al., 2025). This outcome prompted us to pursue the development of more sensitive techniques for molecular sex determination.

By targeting a gene known to have a higher copy number in females (*NPD*), we successfully classified unfed nymphs according to their sex using sensitive qPCR methods, including SYBR Green and fluorescent probe-based assays (Fig 1). This approach builds on the growing genomic resources available for *I. scapularis* (De et al., 2023; Nuss et al., 2023) and expands its uses *to R. microplus* as well, supporting cross-species application of the molecular sexing techniques outlined here. Compared to conventional PCR, qPCR offers several advantages as it is more sensitive, cost-effective, and high throughput, making it a practical choice for routine sex determination. A similar qPCR-based method has been successfully used to determine sex in a variety of organisms, including insects such as lepidopterans (Belousova et al., 2019), soft-shell turtles (Literman et al., 2014), as well as pigs and lizards (Ballester et al., 2013; Rovatsos & Kratochvíl, 2017). These studies demonstrate that qPCR is a reliable and widely applicable tool for sex identification across different species.

Among the molecular methods evaluated, ddPCR provided the most precise and consistent quantification of *NPD* copy number, largely due to its partitioning of template DNA into thousands of droplets, enabling absolute quantification without relying on standard curves. In comparison, qPCR methods were effective but displayed higher variability, most noticeably when template concentration was low. SYBR-based qPCR outperformed probe-based qPCR assays in terms of categorizing nymphal samples molecularly, but both were less sensitive than ddPCR. These findings align with previous studies showing that ddPCR often offers better reproducibility and a stronger tolerance to PCR inhibitors in biological samples (Sedláková et al., 2022). Using two genes instead of one, as we showed for *R. microplus*, led to higher confidence in the qPCR assay, suggesting that combining another gene with *NPD* from our CNV list may result in a more sensitive qPCR assay.

Additionally, we measured the body weight of engorged *I. scapularis* nymphs to determine whether there are sex-based differences in weight post-engorgement, as reported in previous studies (Hu & Rowley, 2000). We attempted to classify the three samples that could not be sexed using molecular methods. According to their body weights, these three samples were classified as two females and one male; however, we could not validate these results as the nymphs died before adult eclosion.

Our work here establishes a foundation for developing a pan-tick sex determination assay. Given that this assay works for both XX: XY and XX: XO systems, it will be interesting to explore whether these markers can be used across additional species. Interestingly, in *R. microplus*, several genes on chromosomes previously annotated as chromosomes 2 and 8 appeared to have variable copy number. This result aligns with a recent karyotyping and genome re-assembly by Tidwell et al., (2024), where previously annotated chromosomes 2 and 8 were identified as the X chromosome.

Future studies should explore larger datasets to test the robustness of these assays. Additional genes may also be identified to develop multiplex assays that improve accuracy and throughput. A deeper examination of the *I. scapularis* sex chromosomes, particularly the male-specific and shared (pseudoautosomal) regions, could aid in identifying new genetic markers.

## Conclusion

The non-lethal molecular method described in this study enables sex determination from a single leg using a female-enriched gene target, *NPD*, via SYBR Green qPCR, probe-based qPCR, and ddPCR. This approach is minimally invasive, scalable, and preserves the viability of the tick for downstream analyses, such as behavioral assays, pathogen acquisition/transmission studies, or colony breeding. Importantly, this method applies to both *I. scapularis* and *R. microplus*, demonstrating its versatility across distantly related tick species. By combining sensitivity, species applicability, and non-lethal sampling, our method provides a valuable tool for developmental, ecological, and vector biology studies where sex identification at early life stages is essential.

## Authors’ contributions

M.G.-N., K.J.M, A.S. and M.P. designed the experiments. M.G.-N supervised the project on *I. scapularis*. K.J.M. and M.P. supervised the *R. microplus* work. I.H. conducted bioinformatics searches for CNV. B.F., A.S., N.H., K.E., and S.B. performed PCR and qPCR assays and analyzed the data. B.F. and J.W. conducted a ddPCR assay. S.B. reared nymphs to adults and took images of molted ticks. M.L. reared the *R. microplus* nymphs and adults. M.G.-N., A.S., and M.P. wrote the first draft of the manuscript, and M.G.-N., A.B.N., M.P., and K.J.M. revised and finalized the text. All authors approved the final version of the manuscript.

## Acknowledgements

We would like to thank Dr. Larissa Martins for feeding tick nymphs on rabbits. This project was funded through National Institutes of Health grants R21AI128393 and R21 AI139778, R21AI176352 and R01AI172943 and a National Science Foundation Grant No. 2019609 to M.G-N, as well as Gates Foundation Grants INV-025721 and INV-045130 to Oxitec Ltd. The conclusions and opinions expressed in this work are those of the author(s) alone and shall not be attributed to the Gates Foundation. Under the grant conditions of the Foundation, a Creative Commons Attribution 4.0 License has already been assigned to the Author Accepted Manuscript version that might arise from this submission.

## Data Accessibility

All data supporting this study are included in the manuscript and supplementary materials.

