## Supplementary data for "Copy Number Variation-Based Molecular Sexing of *Ixodes scapularis* and *Rhipicephalus microplus* Immature Stages Using qPCR and ddPCR Approaches"

^¶^Equal contribution

*Corresponding authors

**Figure Legends**

**Fig S1: Visualization of amplified PCR product on a 1.2% Agarose Gel.** PCR amplification of *EF-1α*, *NPD*, and *RAPGEF2* in extracted adult male (M) and female (F) leg gDNA. The housekeeping gene actin was included as a positive control. The expected product size is ~100-150 bp for all primers, with actin expected to be slightly smaller (~90 bp) than the *EF-1α*, *NPD*, and *RAPGEF2*. For size reference DNA ladder II (100 bp – 1000 bp) was used.

**Fig S2: PCR amplification of the actin housekeeping gene in *Ixodes scapularis* nymph DNA samples (N1-23).** The three unclassified samples all produced clear actin amplicons (N9, N17, N21), indicating good DNA quality. Two classified samples showed minimal amplification; however, they had been successfully classified before degradation occurred. For size reference DNA ladder II (100 bp – 1000 bp) was used.

**Fig S3: Duplex PCR assay for *Ixodes scapularis* nymphs**. Twenty-three nymphs (lanes 1–23) using a published protocol (Ronai et al., 2025) targeting a male-specific genomic region (326 bp) and an autosomal gene (406 bp) was used. Two adult males (M) and one adult female (F) were included as positive controls. Nymphs displaying both the 326-bp and 406-bp bands were classified as male (red letter M), while those showing only the 406-bp band were classified as female. A 100-1000 bp DNA ladder II was used for size reference.

**Fig S4: SYBR-based qPCR for sex determination in the first 12 *I. scapularis* nymph samples using *RAPGEF2* gene.** Most results were conflicting or not classifiable, leading to *RAPGEF2* being excluded from further analysis. The orange bar represents the control value derived from a single adult male or female sample. While some nymph samples could be classified when using one sex (adult M or F) as the control, the same samples then failed to align when the opposite sex was used, resulting in inconclusive data.

**Fig S5: ddPCR results showing *NPD*: *EF-1α* copy number ratios for *I. scapularis* nymphal samples with high variability**. Nymphal samples (n= 23) were analyzed using 1 µL of gDNA per reaction. This setup produced high variability and many of the samples displayed large error bars.

**Fig S6: Histograms of relative copy numbers of genes *RAPGEF2* and *NPD* in unsexed *Rhipicephalus microplus* nymphs.** Two hundred and one unsexed nymphs were tested with a dual probe 09876copy number qPCR assay to determine sex via copy number variation. Both assays produced a roughly bimodal distribution, indicating that these genes likely vary in copy number between males and females.

**Table legends**

**Table S1. Candidate genes exhibiting sex-specific copy number variation in *I. scapularis* Gulia-Nuss genome.** Table shows chromosome, gene ID, fold change, and sex-specific enrichment based on comparisons between male read depth and two female HiFi genomes (Female-1: ~1000 kb, Female-2: ~1800 kb). (*) indicate the genes used in this project, which

**Table S2:** Chromosome, gene ID and location based on genome assembly BIME_Rmic_1.3: GCF_013339725.1, and mean fold change of copy number of genes in adult females relative to male *Rhipicephalus microplus*. Several genes on chromosomes 2 and 8 show higher copy number in females than males. (*) indicate the genes used to sex nymphs in this study (*RAPGEF2* and *NPD*). (**) indicates *EF-1α* which was used as a housekeeping gene.

**Fig S1:**

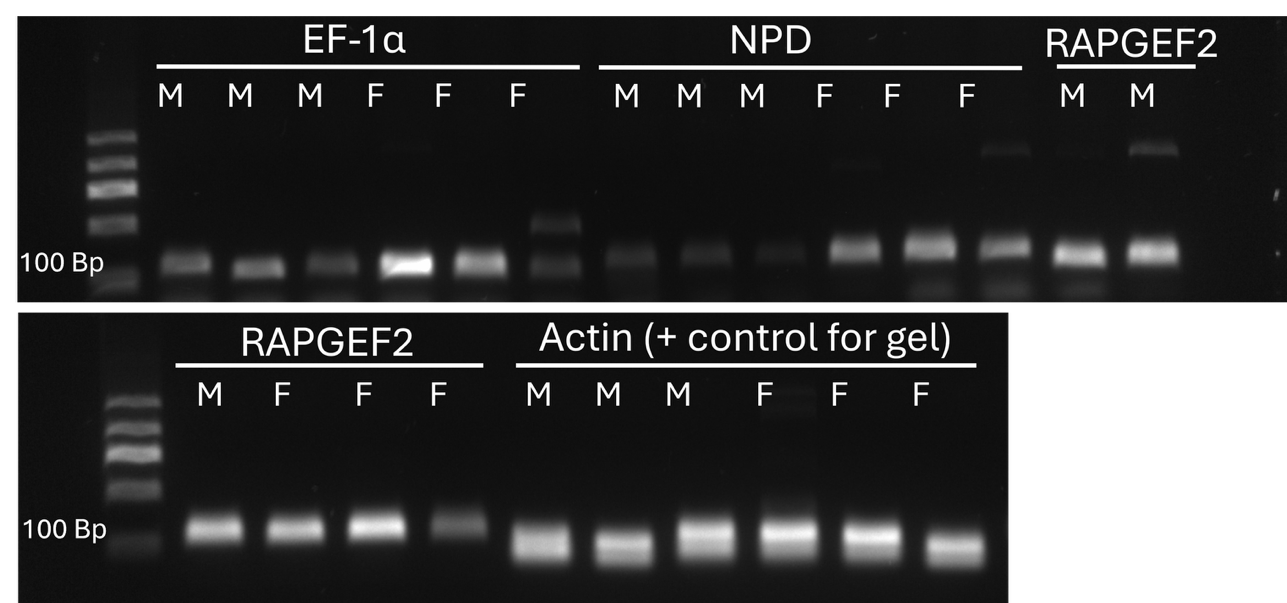

**Fig S2:**

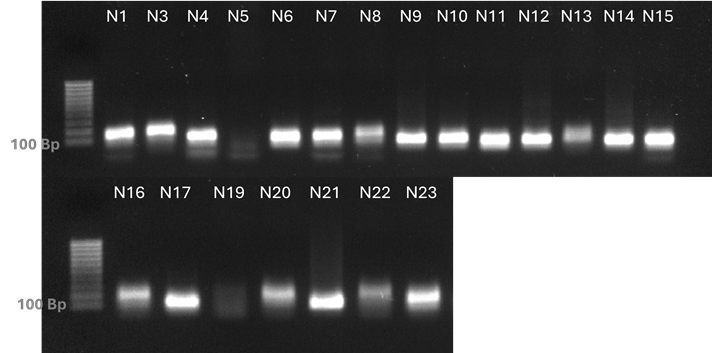

**1000 bp**

**1000 bp**

**Fig S3:**

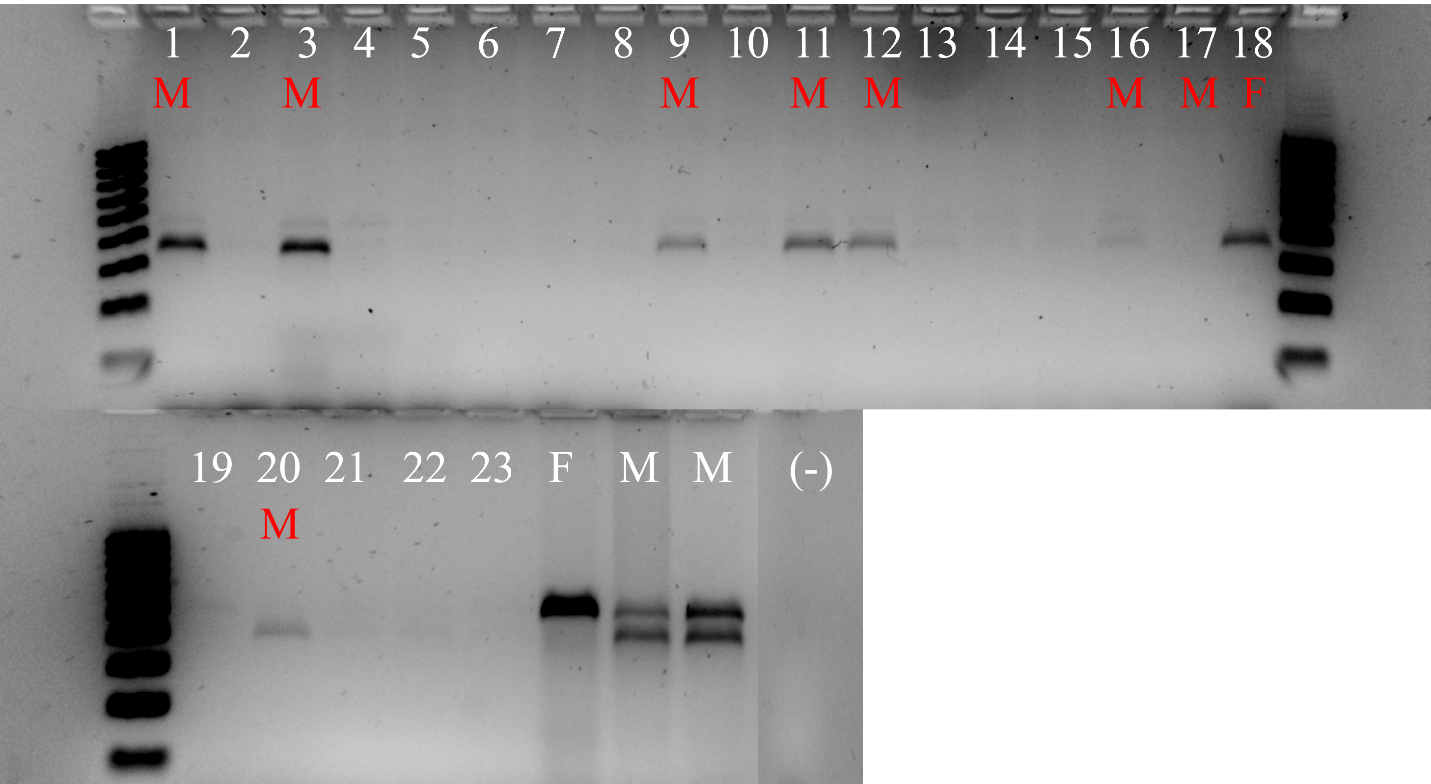

**1000 bp**

**1000 bp**

**100 bp**

**100 bp**

**Fig S4:**

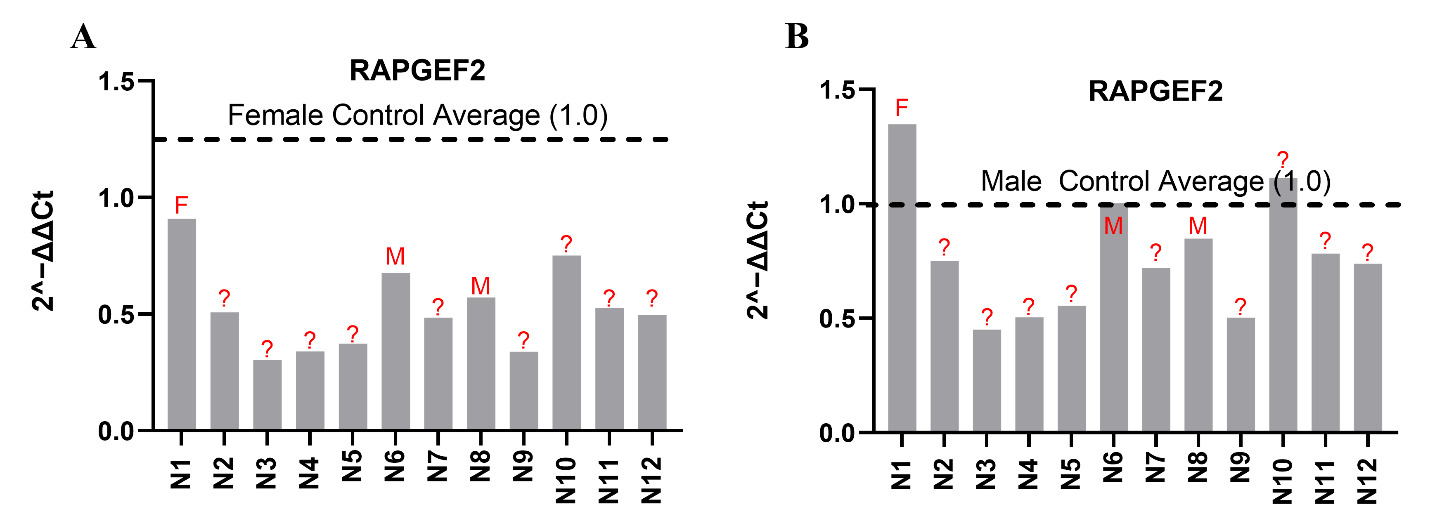

**Fig S5:**

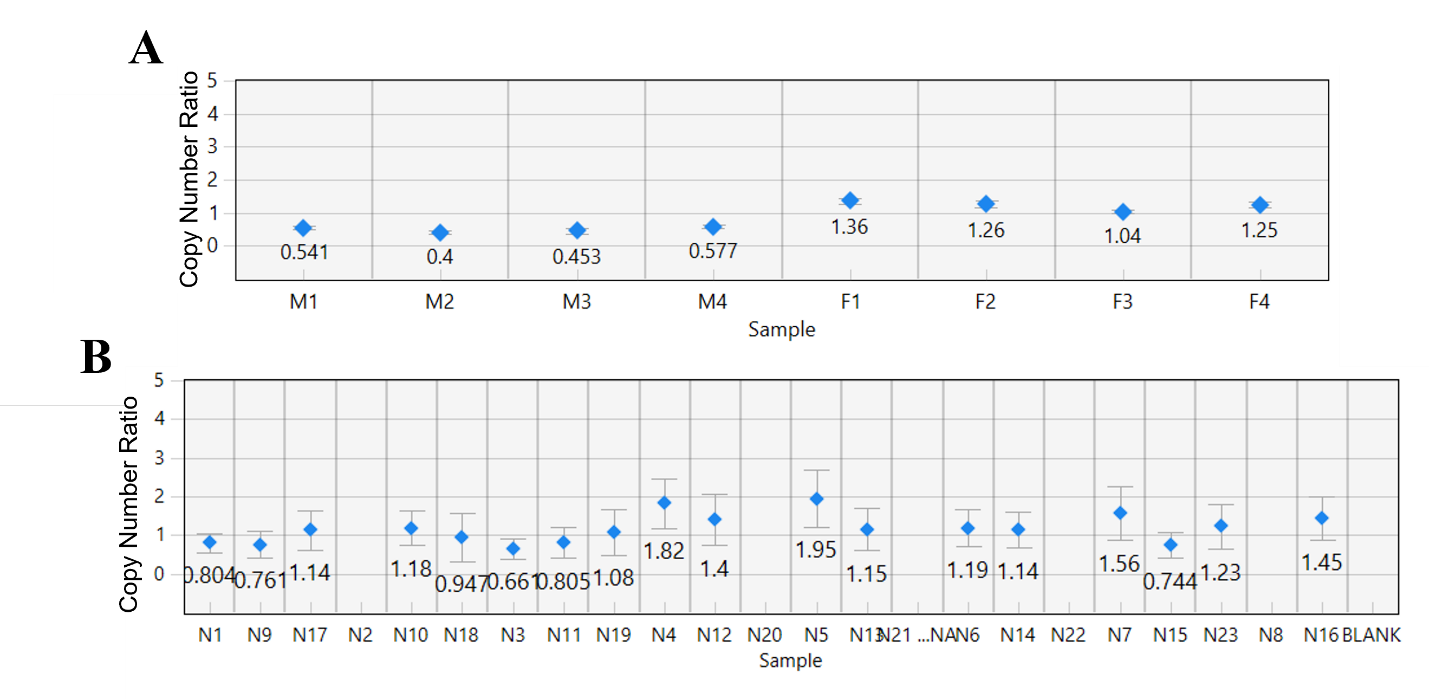

**Fig S6:**

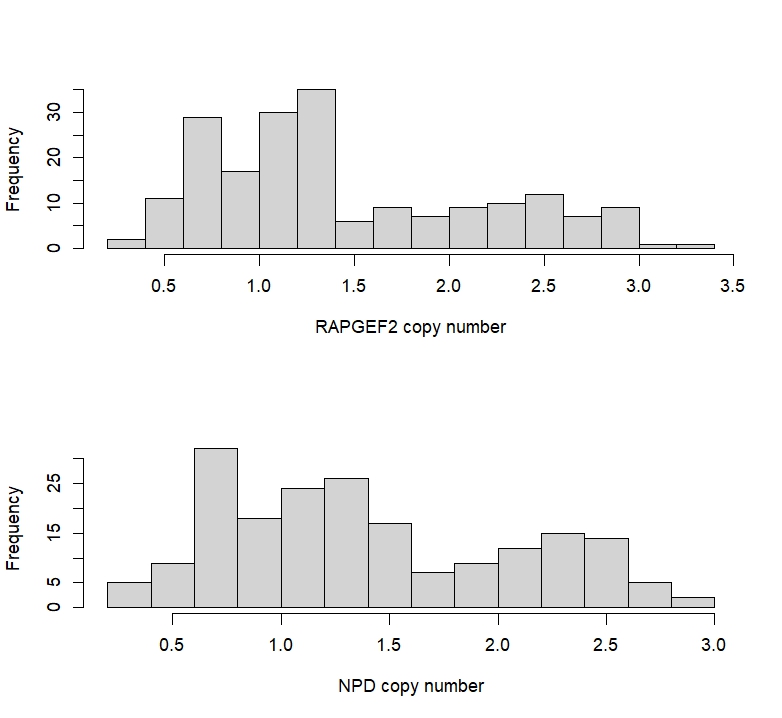

**Table S1**

| **CNV Results**  **Female 1** | | | | | | |
| --- | --- | --- | --- | --- | --- | --- |
| Chromosome | Gene | Start | End | Windows | Fold Change | Enriched in |
| Chr04* | ISCGN121770***** | 224454000 | 224468000 | 14 | 3.7091 | female |
| ChrX | ISCGN302940 | 39780221 | 39913323 | 3 | 2.9562 | female |
| Chr12 | ISCGN279270.1 | 84182973 | 84321307 | 5 | 0.2280 | male |
| Chr12 | ISCGN279350.1 | 84709374 | 84780303 | 12 | 0.2892 | male |
| Chr01 | ISCGN013600.1 | 164356965 | 164418915 | 19 | 0.2390 | male |
| Chr12 | ISCGN271940.1 | 28513034 | 28554606 | 3 | 0.1949 | male |
| **Female 2** | | | | | | |
| Chr04 | ISCGN121770 | 224450273 | 224504838 | 14 | 3.4080 | female |
| ChrX* | ISCGN302940***** | 39780221 | 39913323 | 6 | 2.2527 | female |
| Chr01 | ISCGN024100.1 | 260945948 | 260949532 | 8 | 2.5962 | female |
| Chr12 | ISCGN279360.1 | 84794513 | 84810651 | 4 | 3.1616 | female |
| Chr12 | ISCGN279270.1 | 84182973 | 84321307 | 3 | 1.4128 | female |
| Chr12 | ISCGN279350.1 | 84709374 | 84780303 | 6 | 2.3152 | female |
| Chr01 | ISCGN013600.1 | 164356965 | 164418915 | 3 | 0.4295 | male |
| Chr12 | ISCGN271940.1 | 28513034 | 28554606 | 4 | 0.4310 | male |

**Table S2**

| **Chromosome** | **Gene** | **Start** | **End** | **Mean fold change of copy number in females** |
| --- | --- | --- | --- | --- |
| Chr1 | LOC119186484 | 247321107 | 247324096 | 0.91 |
|  | LOC119186442 | 247053642 | 247076238 | 1.06 |
| Chr2 | LOC119161905 | 83176545 | 83209732 | 1.92 |
|  | LOC119161173 | 124487303 | 124490459 | 1.34 |
|  | LOC119161094* | 101655155 | 102248900 | 1.80 |
| Chr3 | LOC119164383 | 192887113 | 192890725 | 0.99 |
|  | LOC119163128 | 57919511 | 58461070 | 0.89 |
| Chr4 | LOC119167579 | 150246833 | 150252305 | 1.10 |
|  | LOC119167162 | 73155524 | 73162338 | 1.08 |
| Chr5 | LOC119170533** | 70670053 | 70672667 | 1.00 |
|  | LOC119169704 | 96149227 | 96187497 | 0.99 |
| Chr6 | LOC119172232 | 84904124 | 84907889 | 1.13 |
|  | LOC119171436 | 85082314 | 85083740 | 1.39 |
| Chr7 | LOC119173943 | 147745942 | 147747636 | 1.24 |
|  | LOC119174503 | 120647209 | 120658108 | 1.04 |
| Chr8 | LOC119177190 | 146803630 | 146826965 | 1.78 |
|  | LOC119175720 | 20895823 | 20914529 | 1.57 |
|  | LOC119176070* | 58186459 | 58857799 | 2.11 |
| Chr9 | LOC119178352 | 63397086 | 63441064 | 1.13 |
|  | LOC119177784 | 124353430 | 125031430 | 1.01 |
| Chr10 | LOC119179109 | 56502232 | 56870000 | 0.99 |
|  | LOC119179303 | 111264286 | 111525265 | 1.09 |
| Chr11 | LOC119180762 | 74526927 | 74559738 | 1.09 |
|  | LOC119181641 | 69500151 | 69500714 | 0.91 |
